# Genomic characterization of three novel Desulfobacterota classes expand the metabolic and phylogenetic diversity of the Phylum

**DOI:** 10.1101/2021.03.22.436540

**Authors:** Chelsea L. Murphy, James Biggerstaff, Alexis Eichhorn, Essences Ewing, Ryan Shahan, Diana Soriano, Sydney Stewart, Kaitlynn VanMol, Ross Walker, Payton Walters, Mostafa S. Elshahed, Noha H. Youssef

**Author notes:** Corresponding author: Mailing address: Oklahoma State University, Department of Microbiology and Molecular Genetics, 1110 S Innovation Way, Stillwater, OK 74074.

## Abstract

An overwhelming majority of bacterial life remains uncharacterized. Recent efforts to assemble genomes from metagenomes have provided invaluable insights into these yet-uncultured bacterial lineages. We report on the characterization of 30 genomes belonging to three novel classes within the phylum Desulfobacterota. One class (proposed name *Candidatus* “Anaeroferrophillalia”) was characterized by the capacity for heterotrophic growth, either fermentatively or utilizing polysulfide, tetrathionate and thiosulfate as electron acceptors. Autotrophic growth using the Wood Ljungdahl pathway and hydrogen or Fe(II) as an electron donor could also occur in absence of organic carbon sources. The second class (proposed name *Candidatus* “Anaeropigmentia”) was characterized by its capacity for fermentative or aerobic growth at low oxygen thresholds using a broad range of sugars and amino acids, and the capacity to synthesize the methyl/alkyl carrier CoM, an ability that is prevalent in the archaeal but rare in the bacterial domain. Pigmentation is inferred from the capacity for carotenoids (lycopene) production, as well as the occurrence of the majority of genes involved in bacteriochlorophyll *a* biosynthesis. The third class (proposed name *Candidatus* “Zymogenia”) was characterized by the capacity for heterotrophic growth fermentatively using broad sugars and amino acids as carbon sources, and the adaptation of some of its members to hypersaline habitats. Analysis of the distribution pattern of all three classes showed their occurrence as rare community members in multiple habitats, with preferences for anaerobic terrestrial (e.g. hydrocarbon contaminated environments, wetlands, bioreactors), freshwater (e.g. ground water and gas-saturated temperate lakes), and marine (e.g. hydrothermal vents, marine sediments, and coastal sediments) environments, over oxygenated (e.g. pelagic ocean and agricultural land) settings. Special preference for some members of the class *Candidatus* “Zymogenia” to hypersaline environments, e.g. hypersaline microbial mats and lagoons was observed.

**Importance:** Culture-independent diversity surveys conducted in the last three decades have clearly demonstrated that the scope of microbial diversity is much broader than that inferred from isolation efforts. Multiple reasons have been put forth to explain the refractiveness of a wide range of the earth’s microbiome to isolation efforts. Documenting the scope of high-rank phylogenetic diversity on earth, as well as deciphering and documenting the metabolic capacities, physiological preferences, and putative ecological roles of these yet-uncultured lineages represents one of the central goals in current microbial ecology research. Recent efforts to assemble genomes from metagenomes have provided invaluable insights into these yet-uncultured lineages. This study expands our knowledge of the phylum Desulfobacterota through the characterization of 30 genomes belonging to three novel classes. The analyzed genomes were either recovered from Zodletone Spring in southwestern Oklahoma in this study, or recently binned from public metagenomes as part of the Global Earth Microbiome initiative. Our results expand the high-rank diversity within the bacterial tree of life by describing three novel classes within the phylum Desulfobacterota, document the utilization of multiple metabolic processes, e.g. iron-oxidation, aromatic hydrocarbon degradation, reduction of sulfur-cycling intermediates, and features, e.g. coenzyme M biosynthesis, and pigmentation, as salient characteristics in these novel Desulfobacterota classes.

## Introduction

Traditional approaches to characterize bacterial and archaeal taxa have long hinged upon isolation procedures. Despite decades of such efforts, much of the microbial world remains uncultured. Culture-independent surveys utilizing the 16S rRNA gene as a phylogenetic marker have long been utilized to characterize yet-uncultured microbial diversity. However, while useful for documenting the identity, relative abundance, and distribution patterns of microorganisms, such surveys provide limited information on community interactions, metabolic inferences, or microbial growth rates (1–3). The development and wide-scale utilization of genome-resolved metagenomics and single cell genomic approaches have yielded a wealth of metagenome-assembled genome (MAGs) and single-cell amplified genome (SAGs) assemblies (3, 4). In addition to successfully recovering representative genomes of uncultured lineages previously defined by 16S rRNA gene surveys (4), such efforts have also expanded the bacterial tree of life by recovering representatives of novel lineages that previously eluded detection even by culture-independent diversity surveys (5, 6).

Further, the accumulation of MAGs and SAGs has also been leveraged for enacting phylogenomic-based taxonomic schemes that encompass both cultured and uncultured microorganisms (7). The genome taxonomy database (GTDB) utilizes 120 bacterial single-copy genes, average nucleotide identity (ANI), alignment fractionation, and specific quality controls to provide a detailed (phylum to species) genome-based taxonomy (8). The current release (r95) encompasses 111 phyla, 327 classes, 917 orders, 2,282 families, 8,778 genera, and 30,238 species. While broadly agreeing with organismal and 16S rRNA gene-based outlines, e.g. (9), the GTDB proposes several name and rank changes. For example, the Gram-positive Firmicutes are proposed to constitute seven different phyla (Firmicutes A-G). Within the Proteobacteria, the class Betaproteobacteria was reclassified as an order within the class Gammaproteobacteria, and the Epsilonproteobacteria was elevated to a new phylum (Campylobacterota) (7).

Perhaps nowhere was the effect of genomics-based taxonomy more profound than in the class Deltaproteobacteria. The Deltaproteobacteria encompassed anaerobic respiratory and fermentative/syntrophic lineages, the Myxobacteria, *Bdellovibrio*-like predators, and aquatic oligotrophs. The disparate physiological preferences, metabolic capacities, and lifestyles within the Deltaproteobacteria have long been noted, and prior efforts based on single and concatenated gene phylogenies have reported its polyphyletic nature (10). The recent genome-based taxonomy in GTDB r95 has proposed splitting this group into 16 phyla, including 4 distinct cultured phyla: Desulfobacterota, Myxococcota, Bdellovibrionota, and SAR324 (11).

The recently enacted phylum Desulfobacterota encompasses sulfate-reducing and related fermentative and syntrophic lineages previously constituting the bulk of strict anaerobes within the Deltaproteobacteria. GTDB r95 lists 20 classes, 31 orders, 119 families, and 279 genera, of which 12, 14, 38, and 86, respectively, contain cultured representatives. Cultured members of the Desulfobacterota have been identified in a plethora of marine, freshwater, terrestrial, and engineered ecosystems exhibiting wide ranges of salinities, pH, and temperatures (12). Cultured Desulfobacterota show preference to anoxic conditions, and many utilize sulfate, sulfite, thiosulfate, elemental sulfur, or iron (or combinations thereof) as the terminal electron acceptor in respiratory and/or disproportionation processes (13–16). In addition, some grow fermentatively or in syntrophic partnerships (17, 18).

During a broad effort to characterize the yet-uncultured diversity within Zodletone spring, a sulfide and sulfur-rich spring in southwestern Oklahoma, using genome-resolved metagenomics, we recovered multiple MAGs that appear to represent distinct novel classes within the phylum Desulfobacterota. In addition, a recent global effort for binning genomes from publicly available metagenomic datasets (19) yielded additional MAGs belonging to these novel Desulfobacterota classes. Here, we report on the genomic characteristics of 30 MAGs that appear to collectively represent three novel additional classes within the Desulfobacterota. In addition to providing a detailed characterization of their metabolic capacities, physiological preferences, and structural features, we also document their global ecological distribution.

## Materials and Methods

### Sample collection, DNA extraction, and metagenomic sequencing

Samples were collected from the source sediments of Zodletone Spring, located in western Oklahoma’s Anadarko Basin (N34.99562° W98.68895°). The geochemistry of the spring has previously been described (20–23). Samples were obtained from the anoxic sulfidic black sediments at the source of the spring using sterile spatulas and were deposited into sterile 50 mL polypropylene plastic tubes. The samples were transferred to the laboratory on ice and immediately processed. DNA extraction was performed using the DNeasy PowerSoil kit (Qiagen, Valencia, CA, USA) according to manufacturer protocols. The extracted DNA was sequenced using the services of a commercial provider (Novogene, Beijing, China) using the Illumina HiSeq 2500 platform. 281.0 Gbp of raw data were obtained. Contigs were assembled using MegaHit (24), and binned using both Metabat (25) and MaxBin2 (26). DasTool was used to select the highest quality bins from each metagenome assembly (27). Bins that showed contamination levels >5% and/or strain heterogeneity of >10% were further refined and cleaned based on taxonomic affiliations of the bins, GC content, tetranucleotide frequency, and coverage levels using RefineM (28). Bins were classified using the classification workflow option - classify_wf of GTDB-Tk (8) (v 1.3.0), and 5 bins belonging to novel Desulfobacterota classes were selected for further analysis. In addition, we identified genomes belonging to the Desulfobacterota within the recently released 52,515 genomes in the earth microbiome catalogue collection (19) that were deposited in IMG/M database. Of these, 25 genomes belonging to novel Desulfobacterota classes were downloaded from the IMG/M database (May 2020) and included in the analysis.

### Genomes quality assessment and general genomic features

Genome completeness, genome contamination, strain heterogeneity, and GC content were assessed using CheckM (v 1.0.13) (29). Genomes with >50% completion and <10% contamination (n=30) were used for further analysis (Tables S1, S2). Selected MAGs were designated as medium or high-quality drafts based on the criteria set forth by MIMAGs (30). The 5S, 16S, and 23S rRNA sequences were identified using RNAmmer (v 1.2) (31). tRNA sequences were identified and enumerated with tRNAscan-SE (v 2.0.6, May 2020) using the -G general tRNA model (32).

### Phylogenomic analysis

Preliminary classification was carried out using GTDB-Tk (8) with the -classify_wf option. Further phylogenomic analysis was conducted using the 120 single-copy marker genes concatenated alignment that was generated by GTDB-Tk (7). A maximum-likelihood phylogenetic tree was constructed in RAxML (v 8.2.8) (33) with the PROTGAMMABLOSUM62 model and default settings, using members of the phylum Bdellovibrionota as an outgroup. Tree phylogeny, along with average amino acid identity (AAI), calculated using the AAI calculator [http://enve-omics.ce.gatech.edu/]), were used to determine putative taxonomic ranks. The arbitrary AAI cutoffs used were 49%, 52%, 56%, and 68% for class, order, family, and genus, respectively.

### Functional annotation

Protein-coding genes were annotated using Prodigal (v 2.50) (34). Identified protein-coding genes were assigned KEGG orthologies (KO) using BlastKOALA (35), and metabolic pathways were visualized with KEGG mapper (36). For more targeted analysis of functions of interest, hidden markov model (HMM) profile scans were performed on individual genomes. All genomes were queried with custom-built HMM profiles for sulfur metabolism, electron transport chain complexes, and chlorophyll biosynthesis. Custom profiles were built from Uniprot reference sequences for all genes with an assigned KO number, which were downloaded, aligned with Clustal-omega (37), and assembled into a profile with the hmmbuild function of HMMer (v 3.1b2) (38). For genes without a designated KO number, a representative protein was queried against the KEGG genes database using Blastp, and hits with e-values <1e^-80^ were downloaded, aligned, and used to construct an HMM profile as described above. HMM scans were carried out using the hmmscan function of HMMer (38). A thresholding option of -T 100 was used to limit results to alignments with a score of at least 100 to improve specificity. Further confirmation was achieved through phylogenetic assessment and tree building procedures. Briefly, putatively identified sequences were aligned with Clustal-omega (37) against the reference sequences used to build the HMM database and placed into a maximum-likelihood phylogenetic tree using FastTree (v 2.1.10) (39). Sequences that clustered with reference sequences were deemed to be true hits and were assigned a corresponding KO number. FeGenie (40) was used to predict the presence of iron reduction and iron oxidation genes in individual bins. Hydrogenases were identified using HMM scans with profiles constructed from alignments from the Hydrogenase Database (HydDB) (41) using a cutoff e-value of 1e^-20^.

### Search for photosynthetic reaction center

Identification of genes involved in chlorophyll biosynthesis in class “Anaeropigmentia” genomes prompted us to search the genomes for photosynthetic reaction center genes. HMM profiles for Reaction Center Type 1 (RC1; PsaAB), Reaction Center Type 2 (RC2; PufLM and PsbD_1_D_2_) (pfam00223 and pfam00124, respectively), PscABCD (Chlorobia-specific; custom built), PshA/B (Heliobacteria-specific; custom built) (42), as well as the newly identified Psa-like genes from Chloroflexota (43) were used to search the genomes for potential hits using hmmscan. A structurally-informed reaction center alignment (42, 44) was additionally used. The best potential hits were modeled using the SWISS-MODEL homology modeler (45) to check for veracity.

### Ecological distribution

A near-complete 16S rRNA gene from each class was selected as a representative for querying against 16S rRNA databases. Representative sequences were queried against two different databases: 1. The IMG/M 16S rRNA publicly available assembled metagenomes (46), where an e-value threshold of 1e^-10^, percentage similarity ≥90%, and either ≥80% subject length for full-length query sequences or ≥80% query length for non-full-length query sequences criterion were applied, and 2. The GenBank nucleotide (nt) database (accessed November 2020), using a minimum identity threshold of 90%, ≥80% subject length alignment for near full-length query sequences or ≥80% query length for non-full-length query sequences, and a minimum alignment length of 100 bp. Hits meeting the selection criteria were then aligned with 16S rRNA reference gene sequences from each class using Clustal-Omega (37), and the alignment was used to construct maximum-likelihood phylogenetic trees with FastTree (39). The environmental source of hits clustering with the appropriate reference sequences were then classified with a scheme based on the GOLD ecosystem classification scheme (47). Phylogenetic trees were visualized and annotated in iTol (48).

### Sequence and MAG accessions

Metagenomic raw reads for Zodletone sediment are available under SRA accession SRX9813571. Zodletone whole genome shotgun project was submitted to GenBank under Bioproject ID PRJNA690107 and Biosample ID SAMN17269717. The individual assembled MAGs have been deposited at DDBJ/ENA/GenBank under the accession JAFGAM000000000, JAFGAS000000000, JAFGFE000000000, JAFGIX000000000, JAFGSY000000000. The version described in this paper is version JAFGAM010000000, JAFGAS010000000, JAFGFE010000000, JAFGIX010000000, JAFGSY010000000.

## Results

### Three novel classes within the Desulfobacterota

Thirty Desulfobacterota MAGs binned from Zodletone spring sediments and 12 different locations (Table 1, Table S1) clustered into three distinct clades comprising 7, 17, and 6 genomes, that were unaffiliated with any of the 20 currently recognized Desulfobacterota classes in GTDB r95 taxonomic outline (Table 1, Table S1, Figure 1). Average amino acid identity (AAI) and shared gene content (SGC) values between these genomes and representative genomes from all other classes within the Desulfobacterota ranged between 41.63% ± 0.96% – 42.71% ± 1.04% (AAI), and 44.52% ± 4.28% – 51.61% ± 4.37% (SGC), confirming their distinct suggested position as novel classes within the Desulfobacterota phylum (Table S1). Further, the obtained Relative Evolutionary Divergence (RED) values of 0.38 – 0.42 confirmed the distinct class-level designation for all three lineages (Table S1). 16S rRNA gene sequences extracted from representative genomes placed all three groups as members of the “uncultured Deltaproteobacteria” bin within the class Deltaproteobacteria, Phylum Proteobacteria using the RDP-II taxonomic outline (Table S1). In SILVA taxonomic outline release 138.1 (9), these clades were classified as unclassified members of the order Desulfobacterales (clade 1), members of the phylum Sva085 (clade 2), and as uncultured members of the Desulfobacterota phylum (clade 3) (Table S1). We hence propose accommodating these 30 genomes into three distinct classes, for which the following names are proposed based on defining metabolic characteristics predicted from their genomes as described below: *Candidatus* “Anaeroferrophillalia” (order Anaeroferrophillales, family Anaeroferrophillacea), with the MAG assembly 3300022547 (*Anaeroferrophillus wilburensis*) serving as the type material. The name reflects its preference for anaerobic environments and observed capacity to utilize Fe(II) as a supplementary electron donor in absence of organic substrates. *Candidatus* “Anaeropigmentia” (order Anaeropigmentiales, family Anaeropigmentiaceae), with the MAG assembly 3300022855_4 (*Anaeropigmentus antarcticus*) serving as the type material. The name reflects its preference for anaerobic environments and predicted capacity for pigment biosynthesis. *Candidatus* “Zymogenia” (order Zymogeniales, family Zymogeniaceaea), with the MAG assembly Zgenome_24 (*Zymogenus saltonus*) serving as the type material (GenBank assembly accession number JAFGIX000000000). The name reflects its preference for a fermentative mode of metabolism (zymo: Greek for digestion and fermentation). The representative type material MAG was chosen based on the MAG quality deduced from the % completeness (>90%), % contamination (<5%), and the presence of a rRNA operon in the assembly (Table S2). Below, we provide a more detailed analysis of the metabolic capacities, physiological preferences, and ecological distribution of each of these three classes.

**Figure 1.**
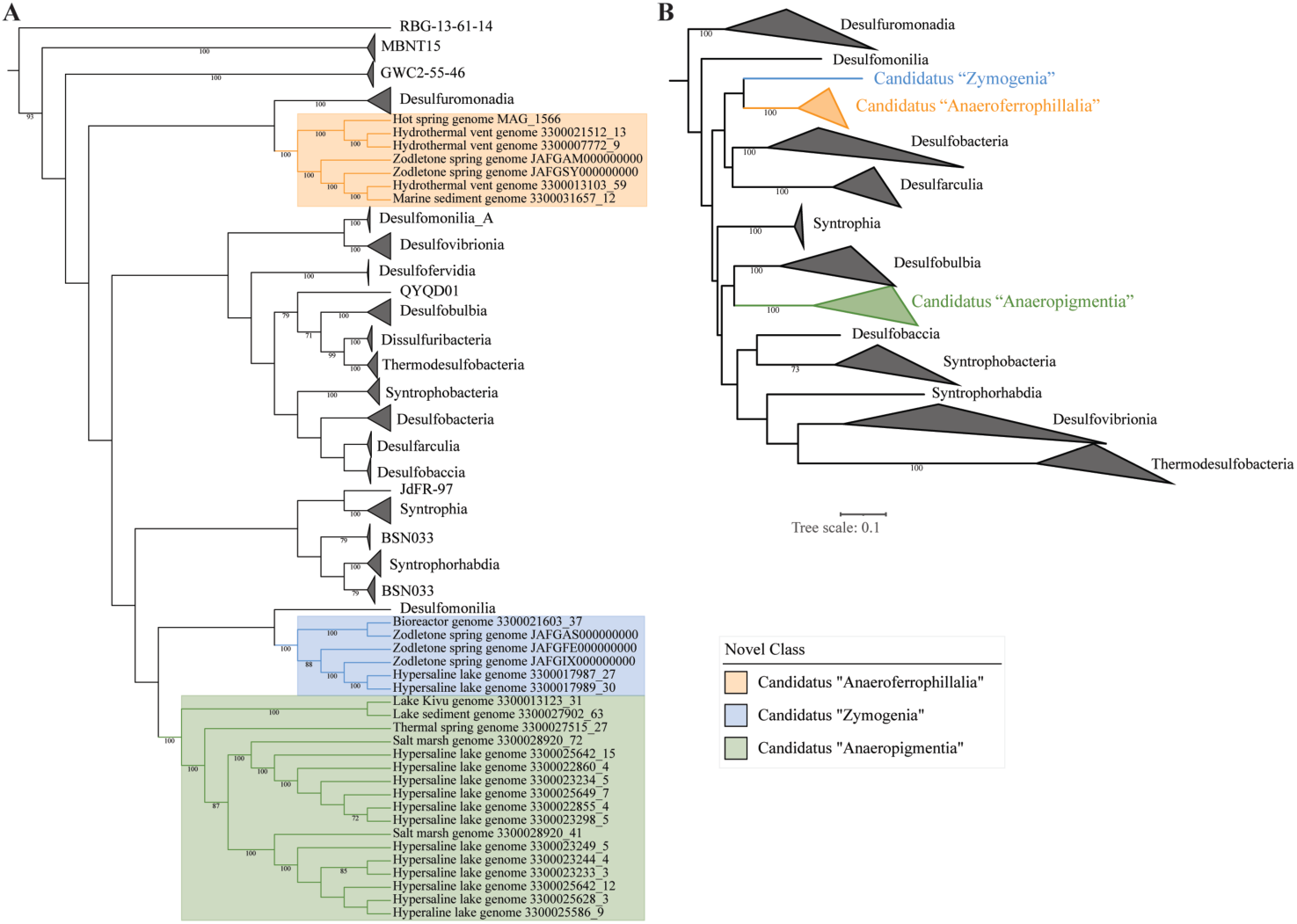
Maximum likelihood phylogenetic tree based on: (A) a concatenated alignment of 120 single-copy genes from all Desulfobacterota classes in GTDB r95, and (B) 16S rRNA gene for Desulfobacterota classes with cultured representatives in GTDB r95. The three novel classes described here are color-coded as shown in the legend. Bootstrap values (from 100 bootstraps) are displayed for branches with ≥70% support. Members of the phylum Bdellovibrionota were used as an outgroup (not shown).

**Table 1.** General genomic features of the studied genomes.

| Bin Name† | GenBank<br>assembly<br>accession<br>number† | Assembly<br>Size<br>(Mbp) | Expected<br>Genome<br>Size (Mbp) | GC % | Number<br>of Genes | Coding<br>% | Ecosystem<br>classification<br>(level 1) | Ecosystem<br>sub<br>classification<br>(level 2) | Site<br>Geographical<br>Location | References |  |
| --- | --- | --- | --- | --- | --- | --- | --- | --- | --- | --- | --- |
|  |  |  |  |  |  |  |  |  |  | NCBI<br>BioProject ID | Citati<br>on |
| 3300007772_9 |  | 1.62 | 2.44 | 48.18 | 1653 | 91.58 | Marine | Hydrothermal<br>vent | Axial<br>Seamount | PRJEB22514 | 78 |
| 3300021512_13 |  | 1.54 | 2.77 | 47.98 | 1543 | 89.89 | Marine | Hydrothermal<br>vent | Guaymas<br>Basin vent | PRJNA444987 |  |
| **3300022547** |  | **2.62** | **2.76** | **54.41** | **2438** | **88.24** | **Freshwater** | **Thermal**<br>**spring** | **Wilbur**<br>**Springs, CA** | PRJNA364781 |  |
| 3300013103_59 |  | 1.56 | 2.91 | 53.95 | 1523 | 89.96 | Marine | Hydrothermal<br>vent | Guaymas<br>Basin vent | PRJNA368391 | 79 |
| 3300031657_12 |  | 1.82 | 2.53 | 47.22 | 1685 | 85.98 | Marine | Sediment | Southern<br>Ocean,<br>Antarctica | PRJNA518182 |  |
| Zgenome_940 | JAFGSY01 | 2.7 | 2.74 | 53.55 | 2471 | 89.72 | Freshwater | Sediment | Zodletone<br>Spring, OK | PRJNA690107 | This<br>study |
| Zgenome_0214 | JAFGAM01 | 2.68 | 3.44 | 63.31 | 2363 | 86.91 | Freshwater | Sediment | Zodletone<br>Spring, OK | PRJNA690107 | This<br>study |
| 3300023233_3 |  | 4.81 | 5.07 | 41.66 | 4638 | 83.06 | Freshwater | Hypersaline | Ace Lake,<br>Antarctica | PRJNA467265 | 80 |
| 3300023244_4 |  | 3.96 | 4.39 | 41.42 | 3755 | 83.06 | Freshwater | Hypersaline | Ace Lake,<br>Antarctica | PRJNA467197 | 80 |
| 3300023249_5 |  | 4.15 | 4.49 | 41.9 | 3893 | 83.31 | Freshwater | Hypersaline | Ace Lake,<br>Antarctica | PRJNA467248 | 80 |
| 3300025586_9 |  | 3.77 | 4.56 | 42.11 | 3529 | 82.98 | Freshwater | Hypersaline | Ace Lake,<br>Antarctica | PRJNA375651 | 80 |
| 3300025628_3 |  | 4.04 | 4.64 | 41.98 | 3788 | 82.85 | Freshwater | Hypersaline | Ace Lake,<br>Antarctica | PRJNA367219 | 80 |
| 3300025642_12 |  | 3.83 | 4.7 | 42.16 | 3578 | 83.09 | Freshwater | Hypersaline | Ace Lake,<br>Antarctica | PRJNA367224 | 80 |
| 3300028920_41 |  | 3.47 | 4.6 | 50.24 | 3129 | 87.61 | Marine | Salt marsh | Falmouth, MA | PRJNA518321 |  |
| **3300022855_4** |  | **3.31** | **3.4** | **48.79** | **2957** | **85.77** | **Freshwater** | **Hypersaline** | **Ace Lake,**<br>**Antarctica** | **PRJNA467247** | **80** |
| 3300023234_5 |  | 3.18 | 3.35 | 49.09 | 2914 | 85.94 | Freshwater | Hypersaline | Ace Lake, Antarctica | PRJNA467206 | 80 |
| 3300023298_5 |  | 3.22 | 3.39 | 48.95 | 2899 | 85.99 | Freshwater | Hypersaline | Ace Lake, Antarctica | PRJNA467209 | 80 |
| 3300025642_15 |  | 3.19 | 3.36 | 48.98 | 2962 | 86.37 | Freshwater | Hypersaline | Ace Lake, Antarctica | PRJNA367224 | 80 |
| 3300025649_7 |  | 3.79 | 3.92 | 48.75 | 3447 | 85.81 | Freshwater | Hypersaline | Ace Lake, Antarctica | PRJNA367218 | 80 |
| 3300028920_72 |  | 2.23 | 3.25 | 57.19 | 2212 | 87.31 | Marine | Salt marsh | Falmouth, MA | PRJNA518321 |  |
| 3300013123_31 |  | 3.2 | 4.73 | 50.1 | 3098 | 84.18 | Freshwater | Temperate | Lake Kivu, Congo | PRJNA404433 |  |
| 3300027902_63 |  | 1.26 | 2.41 | 50.99 | 1326 | 88.93 | Freshwater | Sediment | University of Notre Dame, IN | PRJNA365732 |  |
| 3300027515_27 |  | 2.72 | 3.34 | 50.49 | 2445 | 89.96 | Freshwater | Thermal spring | Thermal spring, YNP | PRJNA367150 |  |
| 3300017987_27 |  | 3.56 | 3.68 | 52.62 | 3284 | 85.75 | Freshwater | Hypersaline | Salton Sea, CA | PRJNA444013 |  |
| 3300017989_30 |  | 3.48 | 3.66 | 52.53 | 3203 | 85.76 | Freshwater | Hypersaline | Salton Sea, CA | PRJNA444014 |  |
| **Zgenome_24** | **JAFGIX01** | **3.76** | **3.84** | **51.37** | **3350** | **84.54** | **Freshwater** | **Sediment** | **Zodletone Spring, OK** | **PRJNA690107** | This study |
| 3300021603_37 |  | 2.2 | 3.54 | 58.54 | 2397 | 87.12 | Engineered | Bioreactor | Toronto, ON | PRJNA501900 |  |
| Zgenome_0311 | JAFGAS01 | 2.94 | 3.65 | 55.99 | 2730 | 86.46 | Freshwater | Sediment | Zodletone Spring, OK | PRJNA690107 | This study |
| Zgenome_1292 | JAFGFE01 | 3.73 | 3.84 | 55.37 | 3398 | 87.75 | Freshwater | Sediment | Zodletone Spring, OK | PRJNA690107 | This study |

†: Accession numbers here include the Metagenome Bin name (numbers starting with 33000; searchable at https://img.jgi.doe.gov/cgi-bin/mer/main.cgi?section=MetagenomeBinSearch&page=searchForm), and the NCBI assembly accession numbers for Zodletone MAGs.
Type strains are listed in **bold**.

### Structural, physiological, and metabolic features

#### Class “Anaeroferrophillalia”

##### General genomic features

Genomes belonging to Class “Anaeroferrophillalia” possess average sized genomes (2.80 ± 0.33 Mbp), GC content (52.66% ± 5.63%), and gene length (937.07 ± 37.07 bp) (Table 1). Structurally, members are predicted to have a Gram-negative cell wall based on the possession of lipopolysaccharide (LPS) biosynthesis-encoding genes, and lack of genes encoding the pentaglycine linkage of peptidoglycan. The presence of the rod-shape determining genes *rodA*/*mreB*, and the genes encoding flagellar assembly suggest rod-shaped flagellated cells. Defense mechanisms include CRISPR defense systems, and Type I restriction endonucleases (Table S3). No evidence for special intracellular structures, e.g. bacterial microcompartments, nanocompartments, or magnetosomes, were identified (Table S3).

##### Physiological features

Members of class “Anaeroferrophillalia” appear to be strict anaerobes, based on the absence of respiratory cytochrome C oxidase (complex IV) components, the presence of the oxygen-limited cytochrome bd complex, and the identification of the oxidative stress enzymes catalase, rubrerythrin, rubredoxin, alkylhydroperoxide reductase, and peroxidase (Table S3). Osmoadaptive capabilities are predicted via the identification of glycine betaine/proline ABC transporter ProXWV.

##### Heterotrophic fermentative capacities

Genomes of Class “Anaeroferrophillalia” possess robust biosynthetic capacities with few amino acids and cofactors auxotrophies (Table S3). The presence of genes encoding the EMP and the non-oxidative PPP pathway indicate heterotrophic growth capabilities. However, a limited number of sugars (glucose, fructose, mannose) appear to support growth (Figure 2A). As well, the capacity to degrade amino acids appears to be limited (Figure 2A, Table S3), and the lack of genes encoding the beta-oxidation pathway precludes potential growth on medium- and long-chain fatty acids. On the other hand, all genomes encode the lactate utilization enzyme D-lactate dehydrogenase (cytochrome) [EC:1.1.2.4], suggesting the capability to grow on D-lactate (Figure 2A). As well, the pathway for anaerobic benzoate metabolism appears to be present in all genomes, suggesting a specialty in degradation of aromatic compounds (Figure 2A). Pyruvate generated from D-lactate or sugar metabolism could be metabolized to acetyl-CoA via pyruvate ferredoxin oxidoreductase [EC: 1.2.7.1] encoded in all genomes, followed by conversion of acetyl-CoA to acetate with concomitant substrate level phosphorylation via the acetate CoA ligase [EC: 6.2.1.13] (Figure 2A, Table S3).

**Figure 2.**
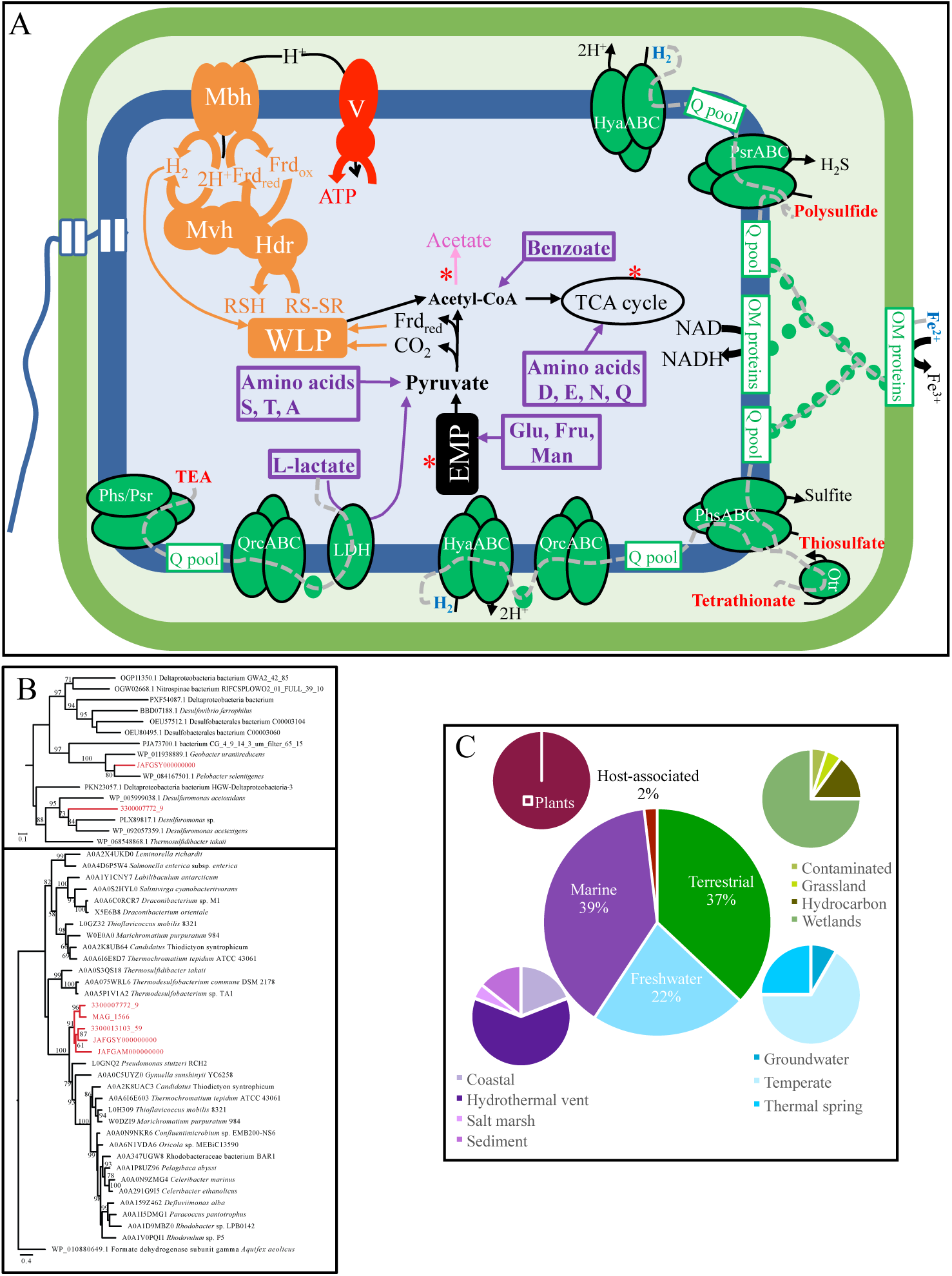
Metabolic reconstruction and ecological distribution for members of the novel class Candidatus “Anaeroferrophillalia”. (A) Cellular metabolic reconstruction based on genomic analysis of 7 genomes belonging to the novel class Candidatus “Anaeroferrophillalia”. Substrates predicted to support growth are shown in purple boxes, electron donors are shown in blue while electron acceptors are shown in red. Fermentation end products are shown in pink. Sites of substrate level phosphorylation are shown as red asterisks. All electron transport chain components in the membrane are shown in green, while components for proton motive force creation and electron carrier recycling are shown in orange. Grey dotted lines depict the predicted flow of electrons from electron donors to electron acceptors. Green spheres in the periplasmic space depict cytochromes. Abbreviations and gene names: EMP, Embden Meyerhoff Paranas pathway; Frd_ox/red_, Ferredoxin (oxidized/ reduced); Fru, fructose; Glu, glucose; Hdr, heterodisulfide reductase complex; HyaABC, periplasmic [Ni Fe] hydrogenase; IM proteins, inner membrane protein complex for the predicted iron oxidation system; LDH, L-lactate dehydrogenase; Man, mannose; Mbh, membrane-­-bound [Ni Fe] hydrogenase; Mvh, Cytoplasmic [Ni Fe] hydrogenase; OM proteins, outer membrane protein complex for the predicted iron oxidation system; Otr, octaheme tetrathionate reductase; PhsABC, thiosulfate reductase; PsrABC, polysulfide reductase; QrcABC, menaquinone reductase; Q pool, quinone pool; RSH/RS-SR, reduced/oxidized disulfide; TCA, tricarboxylic acid cycle; TEA, terminal electron acceptor; V, ATP synthase complex; WLP, Wood Ljungdahl pathway. (B) Phylogenetic affiliation for Candidatus “Anaeroferrophillalia” thiosulfate reductase C subunit (PhsC, top) and the iron oxidation complex protein DFE_0462 (bottom) in relation to reference sequences. Candidatus “Anaeroferrophillalia” sequences are shown in red. Alignments were created in Mafft (76) and maximum likelihood trees were constructed in RaxML (77). Bootstrap support values are based on 100 replicates and are shown for nodes with >50% support. (C) Ecological distribution of Candidatus “Anaeroferrophillalia”-affiliated 16S rRNA sequences. The middle pie chart shows the breakdown of hit sequences based on the classification of the environments from which they were obtained (classification is based on the GOLD ecosystem classification scheme). Further sub-classifications for each environment are shown as smaller pie charts.

##### Respiratory capacities

In addition to fermentative capacities, possible respiratory activities were identified in class “Anaeroferrophillalia”. Possible electron donors identified based on genomic analysis include D-lactate via the D-lactate dehydrogenase [EC: 1.1.2.5]. This enzyme has been studied in several sulfate reducers and its physiological electron acceptor was found to be ferricytochrome c3, which could serve as an entry point to an ETS, with the electrons possibly moving to the genomically-encoded Qrc membrane complex (menaquinone reductase [EC:1.97.-.-], onto the quinone pool and eventually to the terminal electron acceptor. Several hydrogenase-encoding genes were identified in the genomes of class “Anaeroferrophillalia”. These include the periplasmic [Ni Fe] HyaABC (HydDB group 1d), predicted to be involved in hydrogenotrophic respiration (as well as other hydrogenases that are predicted to be involved in recycling reduced equivalents as explained below). Hydrogenotrophic respiration would proceed through the periplasmic hydrogenase moving electrons from H_2_ onto a periplasmic cytochrome C, the Qrc membrane complex, the quinone pool, and eventually to the terminal electron acceptor.

Further, analysis of iron metabolism genes in members of class “Anaeroferrophillalia” genomes indicated their possession of operonic DFE_0448-0451 and DFE_0461-0465 genes, similar to the systems first identified in *Desulfovibrio ferrophilus* (Figure 2B) (49). In the proposed *D. ferrophilus* model, electrons move from an external source, e.g. insoluble minerals like iron, to an outer membrane cytochrome (encoded by DFE_0450 and DFE_0464 by each respective system) through a complex of additional heme-containing periplasmic membrane-bound (DFE_0449 and DFE_0461), periplasmic soluble (DFE_0448 and DFE_0462, DFE_0465), or complex stabilizing (DFE_0451 and DFE_0463) cytochromes. Electrons could potentially then pass on to menaquinone (49) and eventually to the terminal electron acceptor. This later ETS is expected to operate possibly under substrate-limiting conditions (for example in absence of D-lactate) as shown before for *D. ferrophilus* (49).

Possible electron acceptors identified include the sulfur cycle intermediates tetrathionate, based on the identification of genes encoding the octaheme tetrathionate reductase (Otr) (50), as well as the guanylyl molybdenum cofactor-containing tetrathionate reductase (TtrABC) (51). The produced thiosulfate from the action of either Otr or TtrABC could be metabolized through disproportionation, based on the identification of thiosulfate reductase *phsABC* genes (Figure 2B). Polysulfide reduction capability is also predicted based on the identification of genes encoding the membrane-bound molybdoenzyme complex PsrABC (52). No marker genes suggesting the ability to respire sulfate or sulfite were identified. Nitrate reduction genes were similarly lacking.

##### ATP production and recycling reduced equivalents

The genomes encoded complex I components; NADH dehydrogenase [EC: 7.1.12], as well as an F_1_F_o_-ATPase. With an incomplete oxidative phosphorylation pathway, we predict that the NADH dehydrogenase is possibly coupled to the quinone pool and cytochromes for generation of a proton motive force across the inner membrane that can then be used for ATP synthesis via the F_1_F_o_-ATPase, similar to the model predicted in the sulfate reducer *Desulfovibrio vulgaris* (53). Alternatively, or concomitantly, a proton-motive force could possibly be generated during the operation of Wood Ljungdahl (WL) pathway, encoded by all Class “Anaeroferrophillalia” genomes, in the homoacetogenic direction. In that case, a membrane-­-bound mechanism that achieves redox balance between heterotrophic substrate oxidation and the WL function as the electron sink (54, 55) is needed. Candidates for this membrane-bound electron bifurcation mechanism are the membrane-­-bound [Ni Fe] hydrogenase (Mbh) (HydDB group 4d), which couples reduced ferredoxin (produced via the action of pyruvate ferredoxin oxidoreductase [EC: 1.2.7.1]) oxidation to the reduction of protons to H_2_, with the concomitant export of protons to the periplasm (54, 55). Recycling of electron carriers would further be achieved by the cytoplasmic [NiFe]-­-hydrogenase (MvhAGD) plus the heterodisulfide reductase HdrABC, both of which encoded in the genomes (HydDB group [Ni Fe] 3c) (Figure 2A).

##### Autotrophic capacities

In conditions where inorganic compounds, e.g. ferrous ions or H_2_, serve as electron donors, we expect autotrophic capacities to be fulfilled also via the WL pathway. In that case, the proton gradient-­-driven phosphorylation (through the ATPase complex) will be the only means for ATP production (56) as no net ATP gain by substrate level phosphorylation (SLP) is achieved via the WL pathway (55). All genomes encode mechanisms for acetyl-­-CoA (produced from WL pathway) conversion to pyruvate (Pyruvate:ferrodoxin oxidoreductase [EC:1.2.7.1]), reversal of pyruvate kinase (including pyruvate-­-orthophosphate dikinase (EC 2.7.9.1), pyruvate-­-water dikinase [EC 2.7.9.2], as well as pyruvate carboxylase [EC:6.4.1.1] and PEP carboxykinase (ATP) [EC:4.1.1.49]), and the bifunctional fructose-­-1,6-­-bisphosphate aldolase/phosphatase to reverse phosphofructokinase.

#### *Class* “Anaeropigmentia”

##### General genomic features

Members of the Class “Anaeropigmentia” possess relatively large genomes (3.96 ± 0.74 Mbp), with average GC content (47.29% ± 4.55%), and gene length (912.46 ±44.91 bp) (Table 1). Cells are predicted to be Gram-negative rods, with CRISPR systems, type I, and type III restriction endonucleases (Table S3).

##### Physiological features

Class “Anaeropigmentia” genomes encode two distinct pathways for the biosynthesis of the compatible solute trehalose (both from ADP-glucose and glucose via trehalose synthase [EC:2.4.1.245], as well as from UDP-glucose and glucose via the action of trehalose 6-phosphate synthase [EC:2.4.1.15 2.4.1.347] and trehalose 6-phosphate phosphatase [EC:3.1.3.12]), and its degradation via alpha, alpha-trehalose phosphorylase [EC:2.4.1.64]. The genomes also encoded the capability for the storage molecule starch biosynthesis and degradation.

##### Heterotrophic capacities

A heterotrophic lifestyle is predicted, based on the presence of a complete EMP, and ED pathways, a complete TCA cycle, and both the oxidative and non-oxidative branches of the PPP. Substrates predicted to support growth include sugars (glucose, fructose, mannose, sorbitol), amino acids (alanine, aspartate, asparagine, glutamine, glutamate, cysteine, serine), and fatty acids via the beta-oxidation pathway (Table S3, Figure 3A).

**Figure 3.**
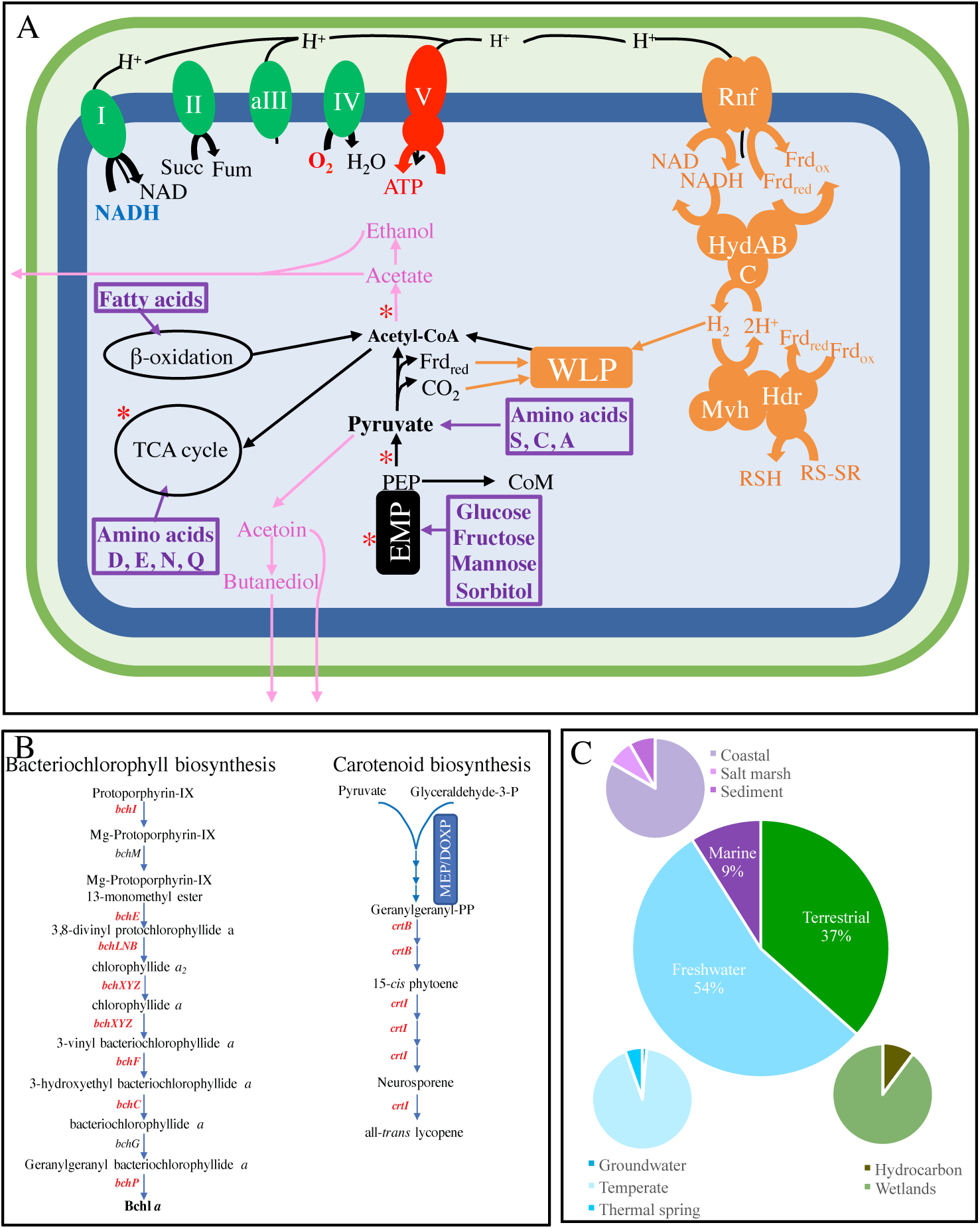
Metabolic reconstruction and ecological distribution for members of the novel class Candidatus “Anaeropigmentia”. (A) Cellular metabolic reconstruction based on genomic analysis of 17 genomes belonging to the novel class Candidatus “Anaeropigmentia”. Substrates predicted to support growth are shown in purple boxes, electron donors are shown in blue while electron acceptors are shown in red. Fermentation end products are shown in pink. Sites of substrate level phosphorylation are shown as red asterisks. All electron transport chain components in the membrane are shown in green, while components for proton motive force creation and electron carrier recycling are shown in orange. Abbreviations and gene names: CoM, coenzyme M; EMP, Embden Meyerhoff Paranas pathway; Frd_ox/red_, Ferredoxin (oxidized/ reduced); fum, fumarate; Hdr, heterodisulfide reductase complex; HydABC, cytoplasmic [Fe Fe] hydrogenase; I, II, aIII, and IV, aerobic respiratory chain comprising complexes I, II, alternate complex III, and complex IV; Mvh, Cytoplasmic [Ni Fe] hydrogenase; PEP, phosphoenol pyruvate; RNF, membrane-bound RNF complex; RSH/RS-SR, reduced/oxidized disulfide; succ, succinate; TCA, tricarboxylic acid cycle; V, ATP synthase complex; WLP, Wood Ljungdahl pathway. (B) Bacteriochlorophylls and carotenoid biosynthesis genes encountered in Candidatus “Anaeropigmentia” genomes. Genes identified in at least one genome are shown in red boldface text, while genes with no homologues in the genomes are shown in black text. MEP/DOXP, the non-mevalonate DOXP/MEP (Deoxyxylulose 5-Phosphate/Methylerythritol 4-Phosphate) pathway for isoprenoid unit biosynthesis. (C) Ecological distribution of Candidatus “Anaeropigmentia”-affiliated 16S rRNA sequences. The middle pie chart shows the breakdown of hit sequences based on the classification of the environments from which they were obtained (classification is based on the GOLD ecosystem classification scheme). Further sub-classifications for each environment are shown as smaller pie charts.

##### Fermentative capacities

The capability of class “Anaeropigmentia” to ferment pyruvate is inferred by the presence of various pathways for end products (butanediol, acetate, ethanol, and acetoin) generation (Table S3, Figure 3A).

##### Respiratory capacities

A complete electron transport chain with complexes I (NADH-quinone oxidoreductase [EC: 7.1.1.2]), II (succinate dehydrogenase [EC: 1.3.5.1]), alternate complex III (encoded by *actABCDEFG)*, and complex IV (cytochrome c oxidase cbb3-type [EC: 7.1.1.9] as well as cytochrome bd ubiquinol oxidase [EC:7.1.1.7]), in addition to a V/A-type as well as F-type H^+^/Na^+^-transporting ATPase [EC: 7.1.2.2], were identified suggesting possible utilization of trace amounts of O_2_ as a terminal electron acceptor by members of Class “Anaeropigmentia”. All Class “Anaeropigmentia” genomes encoded a complete Wood Ljungdahl (WL) pathway. Additional ATP production via oxidative phosphorylation following the generation of a proton-motive force during the operation of WL pathway, is therefore also predicted. In that case, the RNF complex encoded in the majority of genomes would re-­-oxidize reduced ferredoxin at the expense of NAD, with the concomitant export of protons to the periplasm, thus achieving redox balance between heterotrophic substrate oxidation and the WL function as the electron sink (54, 55). Recycling of electron carriers would further be achieved by the cytoplasmic electron bifurcating mechanism (HydABC) plus MvhAGD-­-HdrABC, both of which are encoded in the genomes (HydDB groups [Fe Fe] A3, and [Ni Fe] 3c).

##### Specialized cofactor biosynthesis

Interestingly, the complete pathway encoding the phosphoenol pyruvate-dependent coenzyme M (CoM) biosynthesis was identified in all genomes of Class “Anaeropigmentia” (Figure 3A, Table S3). CoM is a hallmark of methanogenic Archaea, where it acts as a terminal methyl carrier that releases methane upon regeneration of its unmethylated state during methanogenesis (57). However, the utility of CoM in the bacterial domain is less understood. CoM was shown in the bacterial genera *Xanthobacter, Rhodococcus,* and *Mycobacterium* to be involved in propylene degradation as a carrier for a C3 carbon intermediate (58–60). Recently, the bacterial CoM biosynthetic cluster was identified in *X. autotrophicus* Py2 (61). The genes *xcbB1*, *C1*, *D1*, and *E1* encode the bacterial CoM biosynthetic operon, with only *xcbB1* showing homology to the archaeal CoM biosynthesis gene *comA* (61). The remainder of the genes *xcbC1*, *D1*, and *E1* are distinct from the archaeal genes *comBCDE*, and the bacterial biosynthetic pathway proceeds via a different route (61, 62). CoM biosynthesis genes identified in Class “Anaeropigmentia” genomes are distinct from the bacterial CoM biosynthesis genes *xcbC1*, *D1*, and *E1* and are indeed archaeal-like. Searching the functionally annotated bacterial tree of life AnnoTree (63) using the combination of KEGG orthologies corresponding to the archaeal CoM biosynthetic cluster *comABCDE* identified their collective presence in only 14 bacterial genomes from the phyla Acidobacteriota, Actinobacteriota, Bacteroidota, Chloroflexota, Desulfobacterota, Desulfobacterota_B, Latescibacterota, and Proteobacteria. Unfortunately, genes encoding additional enzymes required for propylene degradation (alkene monooxygenase, 2-hydroxypropyl-CoM lyase, 2-hydroxypropyl-CoM dehydrogenase, and 2-oxopropyl-CoM reductase) were absent in all class “Anaeropigmentia” genomes. Therefore, additional research is required to confirm the expression of CoM biosynthesis genes, and further characterize its potential function, if any, in class “Anaeropigmentia”.

##### Pigmentation

The majority of Class “Anaeropigmentia” genomes encode *crtB*, 15-cis-phytoene synthase [EC:2.5.1.32], and *crtI*, phytoene desaturase [EC:1.3.99.26 1.3.99.28 1.3.99.29 1.3.99.31], suggesting the capability of biosynthesis of lycopene from geranylgeranyl-PP. The gene encoding CrtY/L, lycopene beta-cyclase [EC:5.5.1.19], was however missing from all genomes, suggesting an acyclic carotenoid structure (Figure 3B). In addition to carotenoids, a large number of bacteriochlorophyll biosynthesis homologues were encoded in the majority of Class “Anaeropigmentia” genomes. These genes include *bchE* (anaerobic magnesium-protoporphyrin IX monomethyl ester cyclase), *bciB* (3,8-divinyl protochlorophyllide a 8-vinyl-reductase), and *chlLNB* (light-independent protochlorophyllide reductase subunits L, N, and B) catalyzing chlorophyllide a biosynthesis from Mg-protoporphoryin IX. In addition, a near complete pathway for the biosynthesis of bacteriochlolophyll *a* was encoded in the majority of Class “Anaeropigmentia” genomes. This includes the genes *bchXYZ* (3,8-divinyl chlorophyllide a/chlorophyllide a reductase subunits X, Y, and Z), *bchF* (3-vinyl bacteriochlorophyllide hydratase), *bchC* (bacteriochlorophyllide a dehydrogenase), and *chlP* (geranylgeranyl diphosphate/geranylgeranyl-bacteriochlorophyllide a reductase). However, *chlG* (chlorophyll/bacteriochlorophyll a synthase), which catalyzes the penultimate reaction of bacteriochlorophyllide *a* to geranylgeranyl-bacteriochlorophyllide *a*, was not identified (Figure 3B). Despite the presence of the majority of bacteriochlolophyll *a* biosynthetic genes in class “Anaeropigmentia”, all attempts for the identification of a photosynthetic reaction center were unsuccessful.

#### Class “Zymogenia”

##### General genomic features

Genomes belonging to class “Zymogenia” possess average sized genomes (3.7 ± 0.12 Mbp), GC content (54.4% ± 2.7%), and gene length (904.24 ± 62.85 bp) (Table 1). Members of the class “Zymogenia” are Gram-negative rods with CRISPR and Type I restriction endonucleases as defense mechanisms and no intracellular microcompartments (Table S3).

##### Physiological features

Class “Zymogenia” genomes encode the capability for the compatible solute trehalose biosynthesis both from ADP-glucose and glucose via trehalose synthase [EC: 2.4.1.245], as well as from UDP-glucose and glucose via the action of trehalose 6-phosphate synthase [EC:2.4.1.15 2.4.1.347] and trehalose 6-phosphate phosphatase [EC:3.1.3.12]). No genes encoding trehalose degradation were identified in Class “Zymogenia” genomes. Biosynthesis and degradation of the storage molecule starch was encoded in the majority of Class “Zymogenia” genomes. Multiple genes encoding oxidative stress enzymes (catalase, rubrerythrin, rubredoxin, and alkylhydroperoxide reductase, peroxidase, and superoxide reductase) were identified (Table S3).

##### Heterotrophic fermentative capacities

Class “Zymogenia” genomes encode a glycolytic pathway, and a partial TCA pathway, suggesting heterotrophic capacities, possibly on a broad repertoire of sugars including glucose, fructose, galactose, lyxose, arabinose, sorbitol, and xylitol (Figure 4A, Table S3). Surprisingly, genes for fucose degradation to L-lactaldehyde (including L-fucose/D-arabinose isomerase [EC:5.3.1.25, 5.3.1.3], L-fuculokinase [EC:2.7.1.51], and L-fuculose-phosphate aldolase [EC:4.1.2.17]) were encoded in the majority of Class “Zymogenia” genomes, but genes encoding the subsequent conversion of L-lactaldehyde to propanediol (L-lactaldehyde reductase [EC:1.1.1.77]), as well as genes encoding propanediol utilization (propanediol dehydratase [EC: 4.2.1.28]) were missing. Genomes did not encode genes suggestive of aerobic (absence of complex III/ alternate complex III-encoding genes), or anaerobic respiratory capacities, but encoded pyruvate fermentation genes. These include formate C-acetyltransferase [EC: 2.3.1.54] and its activating enzyme [EC: 1.97.1.4] catalyzing pyruvate fermentation to formate and acetyl-CoA, pyruvate ferredoxin oxidoreductase [EC:1.2.7.11] catalyzing pyruvate oxidative decarboxylation to acetyl-CoA, followed by conversion of acetyl-CoA to acetate with concomitant substrate level phosphorylation via the acetate CoA ligase [EC: 6.2.1.13], and acetolactate synthase [EC: 2.2.1.6] and acetolactate decarboxylase [EC:4.1.1.5] for pyruvate conversion to (R)-Acetoin (Table S3, Figure 4A). The lack of a complete electron transport chain, or genes encoding for anaerobic respiratory processes, argue for a predominantly fermentative lifestyle.

**Figure 4.**
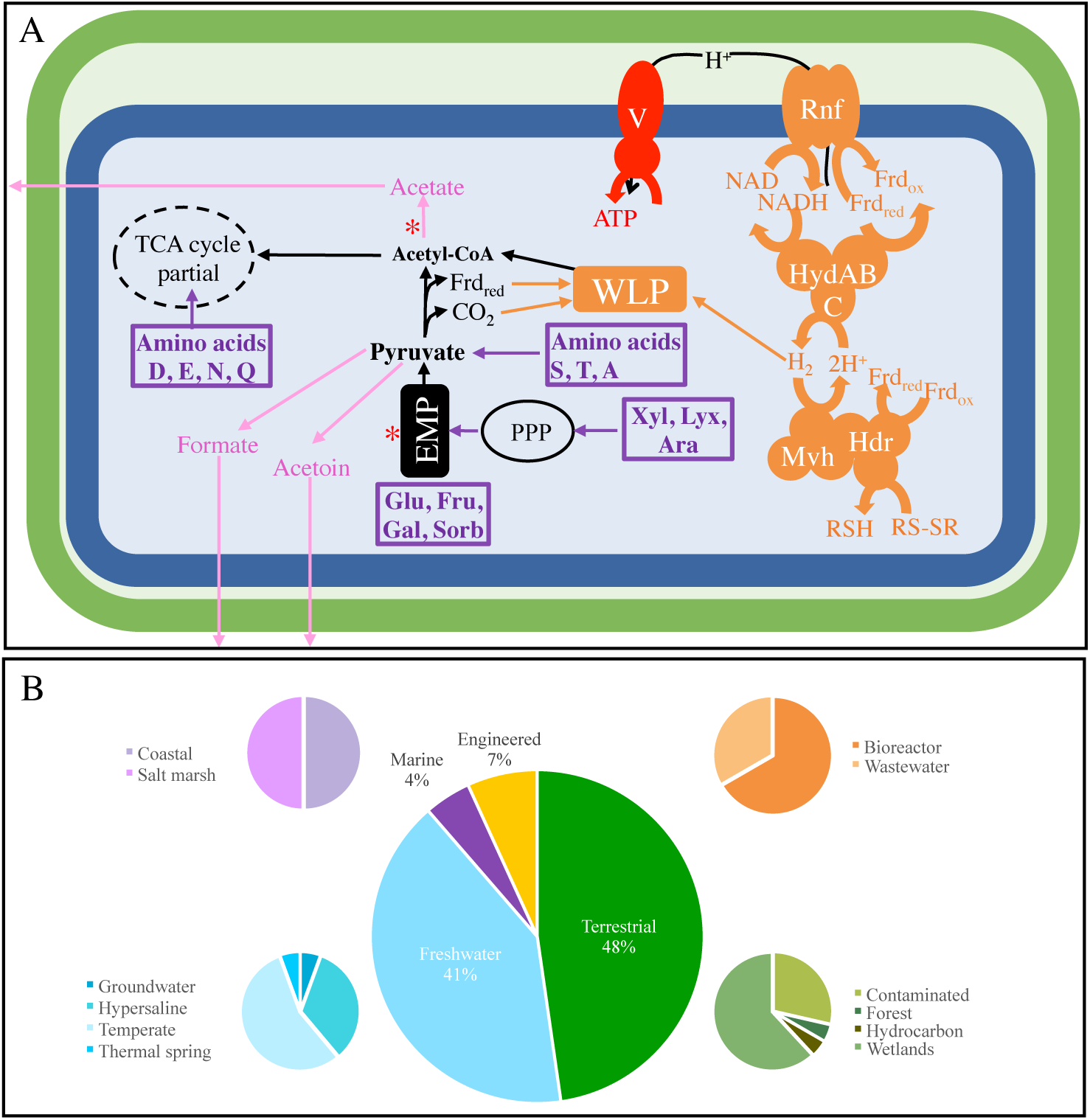
Metabolic reconstruction and ecological distribution for members of the novel class Candidatus “Zymogenia”. (A) Cellular metabolic reconstruction based on genomic analysis of 6 genomes belonging to the novel class Candidatus “Zymogenia”. Substrates predicted to support growth are shown in purple boxes, fermentation end products are shown in pink, and sites of substrate level phosphorylation are shown as red asterisks. Components for proton motive force creation and electron carrier recycling are shown in orange. Abbreviations and gene names: Ara, arabinose; EMP, Embden Meyerhoff Paranas pathway; Frd_ox/red_, Ferredoxin (oxidized/ reduced); Fru, fructose; Gal, galactose; Glu, glucose; Hdr, heterodisulfide reductase complex; HydABC, cytoplasmic [Fe Fe] hydrogenase; Lyx, lyxose; Mvh, Cytoplasmic [Ni Fe] hydrogenase; PPP, pentose phosphate pathway; RNF, membrane-bound RNF complex; RSH/RS-SR, reduced/oxidized disulfide; Sorb, sorbitol; TCA, tricarboxylic acid cycle; V, ATP synthase complex; WLP, Wood Ljungdahl pathway; Xyl, xylitol. (B) Ecological distribution of Candidatus “Zymogenia”-affiliated 16S rRNA sequences. The middle pie chart shows the breakdown of hit sequences based on the classification of the environments from which they were obtained (classification is based on the GOLD ecosystem classification scheme). Further sub-classifications for each environment are shown as smaller pie charts.

##### ATP production and electron carrier recycling

Beside substrate level phosphorylation, ATP production is also possible via the PMF-utilizing H^+^/Na^+^-transporting ATPase [EC: 7.1.2.2] encoded in all genomes. All Class “Zymogenia” genomes encoded a complete Wood Ljungdahl (WL) pathway, and the majority encoded RNF complex. WL pathway in Class “Zymogenia” is predicted to function as an electron sink, and the membrane-bound RNF complex is predicted to help in the generation of a proton motive force across the inner membrane while re-oxidizing the reduced ferredoxin produced from the action of pyruvate ferredoxin oxidoreductase [EC:1.2.7.11]. The PMF generated could be used for ATP production via the complete the H^+^/Na^+^-transporting ATPase [EC: 7.1.2.2]. The cytoplasmic electron bifurcating mechanism (HydABC) plus the heterodisulfide reductase MvhAGD-­-HdrABC (also encoded in the majority of genomes) would function to recycle electron carriers in the cytoplasm (Figure 4A).

#### Ecological distribution

##### Class “Anaeroferrophillalia”

Analysis of ecological distribution pattern identified 54, and 110 16S rRNA gene sequences affiliated with the class “Anaeroferrophillalia” in IMG (Figure 2C), and NCBI nt (Figure S1) databases, respectively (Nov-2020). While “Anaeroferrophillalia” genomes were recovered from a limited number of locations (marine sediments, hydrothermal vents, thermal spring, and Zodletone spring), 16S rRNA analysis expanded their ecological distribution to a range of terrestrial (primarily wetlands and hydrocarbon-impacted environments), marine (predominantly hydrothermal vents, but also coastal and marine sediments), and freshwater (temperate, ground and thermal springs) environments (Figures 2C, S1B-G). The observed distribution patterns reinforce the metabolically predicted preference of members of the “Anaeroferrophillalia” to hypoxic and anoxic settings, as evident by preferences to the oxygen-poor wetlands and hydrocarbon-impacted habitats over grassland and agricultural soils in terrestrial settings, and the preferences to vents and marine sediments over pelagic samples in marine settings. Nevertheless, given the low number of total sequence and the extremely low percentage relative abundance, it is clear that members of the class “Anaeroferrophillalia” are perpetual members of the rare biosphere, and are rarely successful to be a dominant community member in any ecosystem.

##### Class “Anaeropigmentia”

Analysis of ecological distribution pattern identified 134 and 89 16S rRNA gene sequences affiliated with the class “Anaeropigmentia” in IMG (Figures 3C, S1B-G), and NCBI nt (Figure S1A) databases, respectively. While examined genomes were predominantly recovered from hypersaline environments (Ace Lake in Antarctica and Little Sippewissett salt marsh, MA, USA), the majority of 16S rRNA sequences associated with this group were largely associated with non-hypersaline freshwater temperate lake environments, e.g. Yellowstone Lake, the gas-saturated Lake Kivu, and a methane-emitting lake at the University of Notre Dame. Terrestrial environments harboring members of class “Anaeropigmentia” were predominantly wetlands, with hydrocarbon-impacted habitats being the only other terrestrial setting. A limited presence in coastal marine setting and absence from marine pelagic environments was observed. Notably, many of the environments harboring members of class “Anaeropigmentia” are light-exposed, e.g. wetland surface sediment and lake water, justifying pigmentation, although many were not, e.g. coal mine soil and deep lake sediment. Similar to class “Anaeroferrophillalia”, the limited number of affiliated 16S rRNA gene sequences suggests that members of class “Anaeropigmentia” are also part of the rare biosphere, present in small numbers across a range of different habitats.

##### Class “Zymogenia”

Ecological distribution analysis identified only 44 and 33 16S rRNA gene sequences affiliated with the class “Zymogenia” in IMG (Figures 4B), and NCBI nt databases (Figure S1), respectively. A relatively high proportion of class “Zymogenia” 16S rRNA genes were recovered from oxygen-deficient environments, e.g. wetlands, marine sediments, and coastal sediments, as well as aquatic hypersaline settings, e.g. hypersaline lakes (Salton Sea) in California, and lagoons (Etoliko Lagoon in Greece). As well, a significant fraction of these sequences was recovered from anaerobic digestors and bioreactors environments, attesting to the preference of these organisms to anaerobic settings and adaptability of some its members to hypersaline environments (Figures 4B, S1B-G).

## Discussion

Genomic analysis for members of class “Anaeroferrophillalia” revealed the capability of heterotrophic growth on a limited number of substrates, either fermentatively or using sulfur-cycle intermediates (polysulfide, thiosulfate, and tetrathionate) as electron acceptors. Autotrophic growth using the WL pathway and utilization of H_2_ or Fe(II) as electron donors for chemolithotrophic growth in absence of organic carbon sources could also be inferred. Analysis of ecological distribution patterns identified the occurrence of class “Anaeroferrophillalia” as a rare component in few, mostly anaerobic, habitats. Such limited distribution could be a reflection of the limited range of substrates supporting its growth, as well as its dependence on the sulfur-cycle intermediates thiosulfate and tetrathionate as electron acceptors, rather than the more abundant, stable, and ubiquitous sulfate. Although uncommon, microorganisms depending on specific sulfur intermediates (thiosulfate, S, sulfite, tetrathionate), but not sulfate, for growth have previously been reported (64, 65). Such pattern could be a reflection of metabolic interdependencies between various members of the sulfur cycle, a concept formulated based on the identification of a wide range of incomplete pathways in sequenced MAGs (5, 66). In addition, previous studies have shown that thiosulfate and other sulfur cycle intermediates (67, 68), as well as iron (69), were present in much higher levels, and played a more pronounced role in supporting microbial growth on earth during geological eons preceding the evolution of oxygenic photosynthesis. Oxygen production and accumulation from photosynthesis has led to the slow but inexorable oxidation of earth’s surface (great oxidation event), and the establishment of sulfate as the predominant electron acceptor in the sulfur cycle (67). As such, rare lineages depending on H_2_, Fe, and sulfur-intermediates (but not sulfate) for growth could represent lineages that thrived in a preoxygenated earth, but are rare nowadays.

Genomic analysis for members of Class “Anaeropigmentia” revealed a heterotrophic group of microorganisms capable of growing on sugars, amino acids, and fatty acids. Preference appears to be for anaerobic habitats, where it grows fermentatively, with predicted ability to grow under low oxygen microaerophilic conditions. Cells appear to be pigmented with carotenoids (Lycopene). While carotenoid pigments are known to be present in photosynthetic bacteria, where they increase the efficiency of photosynthesis by absorbing in the blue-green region then transferring the absorbed energy to the light-harvesting pigments (70), they also occur in a wide range of non-photosynthetic organisms, including Desulfobacterota, where they serve different purposes, including protection against desiccation (71), radiation (72), and oxidation (73). The occurrence of members of the class “Anaeropigmentia” in transiently and intermittently light-exposed habitats, e.g. lakes and wetlands, could justify their pigmentation, as well as their capability to detoxify trace amounts of oxygen via respiration. More intriguing is the presence of a near-complete machinery for bacteriochlorophyll *a* biosynthesis, a trait exclusive for photosynthetic organisms. However, despite the presence of these biosynthetic pathways, all our attempts for the identification of a photosynthetic reaction center (PRC) were unsuccessful. We argue that the failure to detect PRC-encoding genes in their genomes could be the result of either technical limitations, or the true absence of these genes. Technical limitations include, but are not limited to, contigs harboring PRC-encoding genes not binning from metagenomes; although this is highly unlikely when examining 17 genomes, many of which (n=9) have > 90% completion, and are binned from multiple metagenomic datasets using a variety of approaches. Another possibility for the failure of PRC genes identification could be the high sequence dissimilarity to known reaction center genes. Assuming true absence of PRCs, and hence photosynthetic capacities in class “Anaeropigmentia”, how can the presence and maintenance of most chlorophyll biosynthesis genes be justified? One possibility is the evolution of class “Anaeropigmentia” from a photosynthetic ancestor, with subsequent loss of PRC and some bacteriochlorophyll biosynthesis genes. Indeed, fragmented photosynthetic-related pathways have been previously discovered in multiple genomes, as we described recently (74).

While the loss of the hugely beneficial photosynthetic capacity appears counterintuitive, even implausible, it could be explained in the context of a photosynthetic ancestor evolution to harvest thermal light in an ancient chemolithotrophic world, as previously proposed in (74, 75). These findings demonstrate that bacteriochlorophyll production alone should not be taken as a good measure of photosynthetic capability, and that fragmented pathways must be examined carefully when using bioinformatic tools to prevent any over-assumptions of function.

Finally, our genomic and ecological distribution pattern analysis for members of class “Zymogenia” predicted predominantly anaerobic fermentative organisms; and that some members of this novel class are adapted to growth under saline settings. We also note its extreme rarity in most examined settings. The underlying reason for such rarity is unclear, given its relatively wide substrate utilization range.

In conclusion, our work expands the metabolic and phylogenetic diversity of the Desulfobacterota through the description of 3 novel classes. Our analysis adds to the repertoire of ecological distribution and metabolic capabilities known for the phylum Desulfobacterota, with notable metabolic findings of iron metabolism, thiosulfate and tetrathionate reduction, near complete bacteriocholorophyll biosynthesis, carotenoid biosynthesis, CoM biosynthesis, and fermentation. Ecological distribution patterns observed reinforced and added context to the predicted metabolic capacities gleaned from genomic analysis. The study, overall, demonstrates the utility of bioinformatic tools in exploring and defining unculturable organisms, helping to bridge the vast knowledge gap presented by the uncultured majority.

## Acknowledgments

This work has been supported by NSF grant 2016423 to NHY and MSE. We thank the Hatzenpichler (Montana State University), Edwards (University of Toronto), Bryant (Pennsylvania State University), Biddle (University of Delaware),and Cavicchioli (U of New South Wales), laboratories for permitting access to metagenomes.

